# Green Solvothermal Synthesis of Nitrogen-Doped Chamomile-Derived Carbon Dots with Superior Quantum Yield and Bioimaging Potential: A Comparative Physicochemical Evaluation

**DOI:** 10.64898/2026.05.09.724057

**Authors:** Jagdish Trikam Lagdhir, Satyam Bhalerao, Bhagyesh Parmar, Dhiraj Bhatia

## Abstract

Conventional fluorescent imaging probes, including organic dyes and semiconductor quantum dots, suffer from inherent limitations such as photobleaching, cytotoxicity, poor aqueous dispersibility, and complex synthetic routes, necessitating the development of next-generation nanoscale fluorophores suitable for biological imaging. Carbon dots (CDs) have emerged as a compelling alternative owing to their nanoscale dimensions, tunable photoluminescence, excellent biocompatibility, and amenability to green synthesis from biomass-derived precursors. Herein, we report a comparative synthesis and systematic physicochemical evaluation of nitrogen-doped and undoped carbon dots derived from chamomile (*Matricaria chamomilla* L.) extract, prepared via solvothermal and microwave-assisted routes. Among the four synthesized variants—CM ST-U, CM ST-N, CM MW-U, and CM MW-N—the solvothermally synthesized nitrogen-doped carbon dots (CM ST-N) exhibited markedly superior optical performance, characterized by a high fluorescence quantum yield of 57.2%, which is among the highest reported for biomass-derived nitrogen-doped carbon dots. Comprehensive characterization using UV–visible spectroscopy, photoluminescence (PL) spectroscopy, Fourier-transform infrared (FTIR) spectroscopy, X-ray photoelectron spectroscopy (XPS), dynamic light scattering (DLS), zeta potential analysis, and atomic force microscopy (AFM) confirmed the nanoscale dimensions (~8.3 nm), surface-rich functional groups, successful nitrogen incorporation (10.86 %), and moderate colloidal stability (zeta potential: −17.3 mV). Photoluminescence stability studies across seven solvent systems including biologically relevant media—phosphate-buffered saline (PBS), Dulbecco’s modified Eagle’s medium (DMEM), and serum-free medium (SFM) demonstrated sustained fluorescence emission over 72 hours. In vitro cytotoxicity assessment using the MTT assay on RPE-1 retinal pigment epithelial cells confirmed high cell viability (>70%) across a broad concentration range (10–500 µg mL^−1^) over multiple exposure durations. Collectively, these results establish CM ST-N as a highly fluorescent, biocompatible, and colloidally stable nanoprobe with strong potential for fluorescence-based bioimaging applications.

## 1. Introduction

Fluorescence bioimaging constitutes a cornerstone of contemporary biomedical research, enabling non-invasive visualization of cellular architecture, subcellular dynamics, and molecular interactions with high spatial and temporal resolution.^1,2^ The precision with which fluorescent probes can delineate biological structures at the nanoscale has rendered them indispensable tools in cell biology, cancer diagnostics, drug delivery tracking, and translational medicine.^3,4^ Despite the widespread utility of conventional fluorescent agents, including organic dyes such as fluorescein isothiocyanate (FITC) and rhodamine derivatives, as well as heavy-metal-based semiconductor quantum dots (QDs), their clinical and research translation remains constrained by a series of well-documented physicochemical limitations.^5,6^ Organic dyes are typically characterized by rapid photobleaching under sustained excitation, narrow absorption cross-sections, poor photostability in complex biological matrices, and significant batch-to-batch variability.^7,8^ Semiconductor QDs, while exhibiting broad absorption and narrow emission spectra advantageous for multiplexed imaging, are intrinsically associated with heavy metal cytotoxicity arising from cadmium, lead, or arsenic constituents, raising substantial biosafety concerns that limit their clinical applicability.^9,10^ Consequently, there exists a pressing and unmet need for the development of fluorescent nanoprobes that combine high quantum yield, excellent photostability, robust biocompatibility, aqueous dispersibility, and facile synthesis from sustainable, non-toxic precursors.

Carbon dots (CDs) have attracted considerable scientific interest over the past two decades as a versatile class of zero-dimensional carbon-based nanomaterials, typically sub-10 nm in diameter, exhibiting intrinsic photoluminescent properties.^11,12^ First observed incidentally during the electrophoretic purification of single-walled carbon nanotubes in 2004, CDs have since been established as a structurally diverse family encompassing graphene quantum dots (GQDs), carbon quantum dots (CQDs), carbon nanodots (CNDs), and carbonized polymer dots (CPDs), each distinguished by their formation mechanism, degree of graphitization, and surface chemical landscape.^13,14^ Among the properties that render CDs particularly attractive for bioimaging, their excitation-dependent photoluminescence, high resistance to photobleaching relative to organic dyes, intrinsic aqueous dispersibility conferred by surface oxygen-containing functional groups, and low cytotoxicity are most consequential.^15,16^ The optical behavior of CDs is governed by a combination of quantum confinement effects, surface defect states, and the chemical nature of heteroatom-containing functional groups, all of which are modulated by synthetic conditions and precursor composition.^17,18^

Heteroatom doping, particularly nitrogen incorporation, has emerged as a widely adopted and effective strategy for enhancing the photoluminescence efficiency of carbon dots.^19,20^ The substitution of carbon atoms with nitrogen within the sp^2^-hybridized graphitic framework, or the introduction of nitrogen-containing surface functional groups such as amino (–NH_2_) and amide (–CONH–) moieties, modifies the electronic band structure of CDs, creating additional emissive surface states and narrowing the effective bandgap.^21,22^ These electronic modifications collectively enhance radiative recombination efficiency and quantum yield, both of which are critical determinants of fluorescence probe performance in bioimaging applications.^23,24^ Nitrogen-doped CDs synthesized via hydrothermal and solvothermal routes have consistently demonstrated quantum yields substantially exceeding those of their undoped counterparts, with values ranging from 15% to over 60% reported across different precursor systems.^25,26^

In parallel, the imperative to align nanomaterial synthesis with the principles of green chemistry has driven increasing interest in biomass-derived carbon precursors as sustainable alternatives to reagent-grade organic molecules.^27,28^ Naturally occurring plant extracts, agricultural residues, and food-grade materials are rich in carbon, oxygen, and nitrogen-containing biomolecules—including polyphenols, flavonoids, organic acids, and amino acids— that serve simultaneously as carbon sources, natural passivating agents, and intrinsic nitrogen dopants during carbonization.^29,30^ Carbon dots derived from sources such as citric acid, orange peel, banana peel, lemon juice, and various medicinal plant extracts have been reported to exhibit strong fluorescence, good biocompatibility, and favorable surface chemistry without recourse to toxic reagents or complex multi-step synthesis.^31,32^ Solvothermal and hydrothermal methods are particularly well-suited to green CD synthesis, as they operate under closed-system conditions, allow precise control over reaction temperature and duration, promote uniform carbonization, and yield particles with consistent size distributions and well-defined surface chemistries.^33,34^

German chamomile (*Matricaria chamomilla* L.) is a medicinally significant plant whose phytochemical profile includes flavonoids, phenolic acids, terpenoids, and polysaccharides, rendering it a rich and chemically diverse carbon precursor for green CD synthesis.^35,36^ Despite its widespread utilization in the pharmaceutical and nutraceutical industries, chamomile biomass has remained comparatively underexplored as a raw material for the synthesis of functional carbon nanomaterials.^37^ The few studies that have investigated chamomile-derived CDs have predominantly focused on their sensing applications, particularly for the detection of antibiotics and metal ions in food and environmental matrices, with limited systematic investigation of their bioimaging potential.^38,39^ Furthermore, comparative studies evaluating the influence of synthesis route—specifically solvothermal versus microwave-assisted methods—and nitrogen doping on the physicochemical and optical properties of chamomile-derived CDs remain absent from the literature, representing a critical knowledge gap.

Bioimaging applications impose stringent requirements on fluorescent nanoprobes: high quantum yield to enable sensitive detection at low probe concentrations, nanoscale dimensions to facilitate cellular internalization, photoluminescence stability in physiologically relevant media, and cytotoxic safety profiles compatible with living cellular systems.^40,41^ The intersection of green synthesis, nitrogen doping, and bioimaging-targeted evaluation represents a compelling and underexplored research direction for chamomile-derived carbon nanomaterials. We earlier shoed developed carbon-based nanomaterials for bioimaging and theranostic applications, demonstrating the utility of carbon quantum dots in cellular imaging, tissue-specific uptake, and endocytic pathway analysis.^4,25,26,34^Building upon this foundational work, the present study addresses the identified research gap through a systematic comparative synthesis and evaluation of nitrogen-doped and undoped chamomile-derived CDs prepared by solvothermal and microwave-assisted routes, with bioimaging applicability as the primary evaluative framework.

Herein, four carbon dot variants-CM ST-U (solvothermal, undoped), CM ST-N (solvothermal, nitrogen-doped), CM MW-U (microwave-assisted, undoped), and CM MW-N (microwave-assisted, nitrogen-doped) were synthesized from chamomile extract and comparatively evaluated using UV–visible spectroscopy, photoluminescence spectroscopy, and FTIR spectroscopy to identify the optimal fluorescent candidate. The superior variant, CM ST-N, was subsequently subjected to comprehensive physicochemical characterization encompassing XPS, DLS, zeta potential analysis, AFM, and quantum yield determination, alongside multi-media photoluminescence stability assessment and in vitro MTT cytotoxicity evaluation. The results demonstrate that solvothermal nitrogen doping of chamomile-derived CDs yields nanoprobes with a quantum yield of 57.2%, photostability across biologically relevant media, nanoscale dimensions favorable for cellular uptake, and excellent biocompatibility, collectively establishing their suitability for fluorescence bioimaging applications.

## 2. Materials and Methods

### 2.1 Chemicals and Reagents

Chamomile liquid extract was sourced from BRM Chemicals (India) and used without further modification. Urea (SRL Chemicals, India) served as the nitrogen dopant for the synthesis of nitrogen-doped variants. Absolute ethanol and methanol were employed as co-solvents during synthesis and characterization. Milli-Q water (18.2 MΩ cm, Merck Millipore) was used throughout all synthesis and characterization procedures. Acids employed for glassware cleaning included concentrated nitric acid (HNO_3_), hydrochloric acid (HCl), and sulfuric acid (H_2_SO_4_). Quinine sulfate dissolved in 0.1 M H_2_SO_4_ served as the reference standard for quantum yield determination. For cytotoxicity studies, Dulbecco’s modified Eagle’s medium (DMEM, Gibco), fetal bovine serum (FBS, Gibco), and the MTT reagent (3-(4,5-dimethylthiazol-2-yl)-2,5-diphenyltetrazolium bromide) were used. Dimethyl sulfoxide (DMSO, HiMedia) was used to dissolve formazan crystals. Phosphate-buffered saline (PBS) was used as the physiological buffer medium. All reagents were of analytical grade and used without additional purification unless otherwise stated.

### 2.2 Synthesis of Carbon Dots

Prior to synthesis, all Teflon-lined hydrothermal vessels and glassware were thoroughly cleaned using freshly prepared aqua regia (HCl:HNO_3_, 3:1 v/v) for 12–24 hours under a fume hood, followed by sequential rinsing with tap water, detergent solution, acetone, and Milli-Q water. All vessels were air-dried completely before use.

#### Solvothermal synthesis of undoped carbon dots (CM ST-U)

A reaction mixture comprising 1 mL chamomile extract, 10 mL Milli-Q water, and 10 mL absolute ethanol (total volume: 20 mL) was prepared in a 50 mL Falcon tube. The mixture was vortexed briefly and subjected to ultrasonication for 10 minutes at approximately 22°C to achieve a homogeneous dispersion. The mixture was subsequently transferred to a Teflon-lined stainless-steel hydrothermal autoclave, sealed, and placed in a muffle furnace at 140°C for 8 hours. After the reaction was complete, the system was allowed to cool naturally to room temperature, followed by cooling under running tap water. The resulting pale yellow-to-light-brown dispersion was filtered sequentially through 0.45 µm and 0.22 µm nylon membrane syringe filters (Ran Disc) to remove larger particulates, yielding a clear CD dispersion stored at 4°C in the dark.

#### Solvothermal synthesis of nitrogen-doped carbon dots (CM ST-N)

The synthesis was performed identically to CM ST-U, with the addition of 100 mg of urea as the nitrogen source, dissolved in the reaction mixture prior to sonication. The autoclave was subjected to thermal treatment at 140°C for 8 hours. The resulting product was purified by sequential membrane filtration (0.45 µm followed by 0.22 µm) and stored at 4°C protected from light.

#### Microwave-assisted synthesis of undoped carbon dots (CM MW-U)

A mixture of 1 mL chamomile extract, 5 mL Milli-Q water, and 5 mL absolute ethanol was prepared in a 15 mL Falcon tube, vortexed, and sonicated for 10 minutes. The homogeneous mixture was transferred to a 200 mL glass beaker and subjected to microwave irradiation at 500 W in repeated 30-second cycles (Whirlpool domestic microwave), with manual agitation between cycles, until complete solvent evaporation and formation of a brownish residue. The cooled residue was reconstituted in 5 mL Milli-Q water and filtered through 0.45 µm and 0.22 µm membrane filters before storage.

#### Microwave-assisted synthesis of nitrogen-doped carbon dots (CM MW-N)

The procedure followed that of CM MW-U with the addition of 50 mg of urea to the reaction mixture prior to microwave treatment. Purification and storage conditions were identical to those described for CM MW-U.

### 2.3 Physicochemical Characterization

UV–visible absorption spectra of all four CD variants were acquired using a UV–visible spectrophotometer over a wavelength range of 200–800 nm using quartz cuvettes. Samples were appropriately diluted in the respective synthesis solvent prior to measurement. Photoluminescence (PL) emission spectra were recorded using a fluorescence spectrophotometer at multiple excitation wavelengths (220–450 nm in 50 nm increments) to evaluate excitation-dependent fluorescence behavior. All samples were measured under identical instrumental settings to permit direct comparative analysis.

FTIR spectroscopy was performed using an ATR-FTIR spectrometer over the range 4000–400 cm^−1^ to identify surface functional groups. XPS analysis was conducted using a Thermo Scientific K-Alpha instrument under ultra-high vacuum conditions. Survey spectra were acquired to determine elemental composition, with C 1s, N 1s, and O 1s binding energy signals used for elemental quantification and chemical state assignment. Samples were prepared by drop-casting diluted dispersions onto clean glass coverslips and drying in a desiccator overnight.

Hydrodynamic size distribution and zeta potential of CM ST-N were measured by dynamic light scattering (DLS) using a Malvern Analytical Zetasizer Nano ZS instrument. Prior to measurement, samples were sonicated briefly and centrifuged to remove aggregates, and only the supernatant was analyzed. Surface morphology and particle distribution were examined by atomic force microscopy (AFM) using a Bruker Nanowizard Sense Plus Bio-AFM in tapping mode under ambient conditions. Diluted CM ST-N dispersion was drop-cast onto freshly cleaved mica and dried in a vacuum desiccator overnight prior to imaging.

### 2.4 Quantum Yield Determination

The fluorescence quantum yield (Φ) of CM ST-N was determined using the comparative method with quinine sulfate in 0.1 M H_2_SO_4_ (Φ*ref* = 0.54) as the reference standard.^42^ Serial dilutions of both quinine sulfate and CM ST-N were prepared to yield absorbance values in the range of 0.04–0.07 at the excitation wavelength of 400 nm, ensuring operation within the linear optical regime and minimizing inner filter effects. PL emission spectra were recorded under identical instrumental conditions, and the integrated emission intensities were calculated from the area under each emission curve. Quantum yield was calculated using the standard relative equation accounting for the refractive indices of the respective solvents.

### 2.5 Photoluminescence Stability Assessment

The photoluminescence stability of CM ST-N was evaluated in seven solvent and biological media systems: Milli-Q water, methanol, absolute ethanol, 50% ethanol, PBS, DMEM, and serum-free medium (SFM). Emission intensity at 400 nm excitation was recorded at 24, 48, and 72 hours under ambient storage conditions. All measurements were performed in triplicate. Relative fluorescence intensity was normalized to the initial reading to enable cross-media comparison.

### 2.6 In Vitro Cytotoxicity Assessment

The in vitro cytocompatibility of CM ST-N was evaluated using the MTT assay on RPE-1 retinal pigment epithelial cells, a non-cancerous human cell line relevant to biocompati ility screening.^43^ Cells were maintained in DMEM supplemented with 10% FBS and 1% penicillin-streptomycin at 37°C in a humidified 5% CO_2_ atmosphere. For cytotoxicity evaluation, 1–2 × 10^4^ cells per well were seeded in 96-well plates and allowed to adhere overnight. Following adhesion, cells were exposed to CM ST-N at concentrations of 10, 50, 100, 250, and 500 µg mL^−1^ for 12, 24, and 48 hours. After each treatment period, medium was aspirated, MTT reagent (5 mg mL^−1^) was added, and cells were incubated for 3–4 hours at 37°C. Formazan crystals were dissolved in DMSO, and absorbance was recorded at 570 nm using a microplate reader. Cell viability was expressed as a percentage relative to untreated controls, calculated using the standard formula:

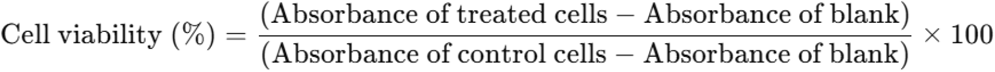

### 2.7 Statistical Analysis

All data are expressed as mean ± standard deviation (SD) from at least three independent experiments. Statistical significance was assessed using one-way analysis of variance (ANOVA) followed by appropriate post-hoc tests using GraphPad Prism software (version 8.0.2). A p-value of less than 0.05 was considered statistically significant.

## 3. Results and Discussion

### 3.1 Synthesis and Preliminary Optical Screening

Four carbon dot variants were synthesized from chamomile extract using two routes— solvothermal and microwave-assisted—each applied with and without urea as a nitrogen dopant. The solvothermal route employed a Teflon-lined autoclave at 140°C for 8 hours, conditions that promote controlled carbonization and uniform particle formation through sustained thermal treatment under autogenous pressure.^29,33^ In contrast, the microwave-assisted route achieved rapid solvent evaporation through dielectric heating at 500 W in 30-second cycles, offering a kinetically driven alternative that typically produces particles with broader size distributions and less-developed graphitic domains due to the comparatively shorter and less uniform heating profile.^44,45^ Upon UV illumination, all four samples exhibited visible fluorescence emission, confirming the formation of photoluminescent carbon-based nanostructures from chamomile biomass. However, marked differences in emission intensity were immediately apparent, with solvothermally prepared samples demonstrating considerably brighter fluorescence than their microwave-derived counterparts, and nitrogen-doped variants outperforming their undoped equivalents under identical conditions. These preliminary qualitative observations guided the selection of CM ST-N as the primary candidate for detailed characterization and bioimaging-targeted evaluation.

### 3.2 UV–Visible Spectroscopy

UV–visible absorption spectra of all four synthesized variants exhibited a characteristic strong absorption band in the range of 218–226 nm (Fig. 2), attributable to π→π* electronic transitions arising from C=C bonds within the sp^2^-hybridized graphitic carbon core, a spectral feature commonly associated with the formation of carbonaceous nanostructures.^39,46^ A secondary, weaker absorption shoulder was observed at longer wavelengths in all samples, consistent with n→π* transitions associated with C=O and other surface oxygen-containing functional groups introduced during the carbonization of chamomile biomass.^17,47^ The presence of this secondary band across all variants confirms that surface oxidation occurred during synthesis, resulting in the incorporation of carboxyl, carbonyl, and hydroxyl moieties that are functionally relevant for aqueous dispersibility and surface passivation critical to bioimaging performance.

**Figure. 1.**
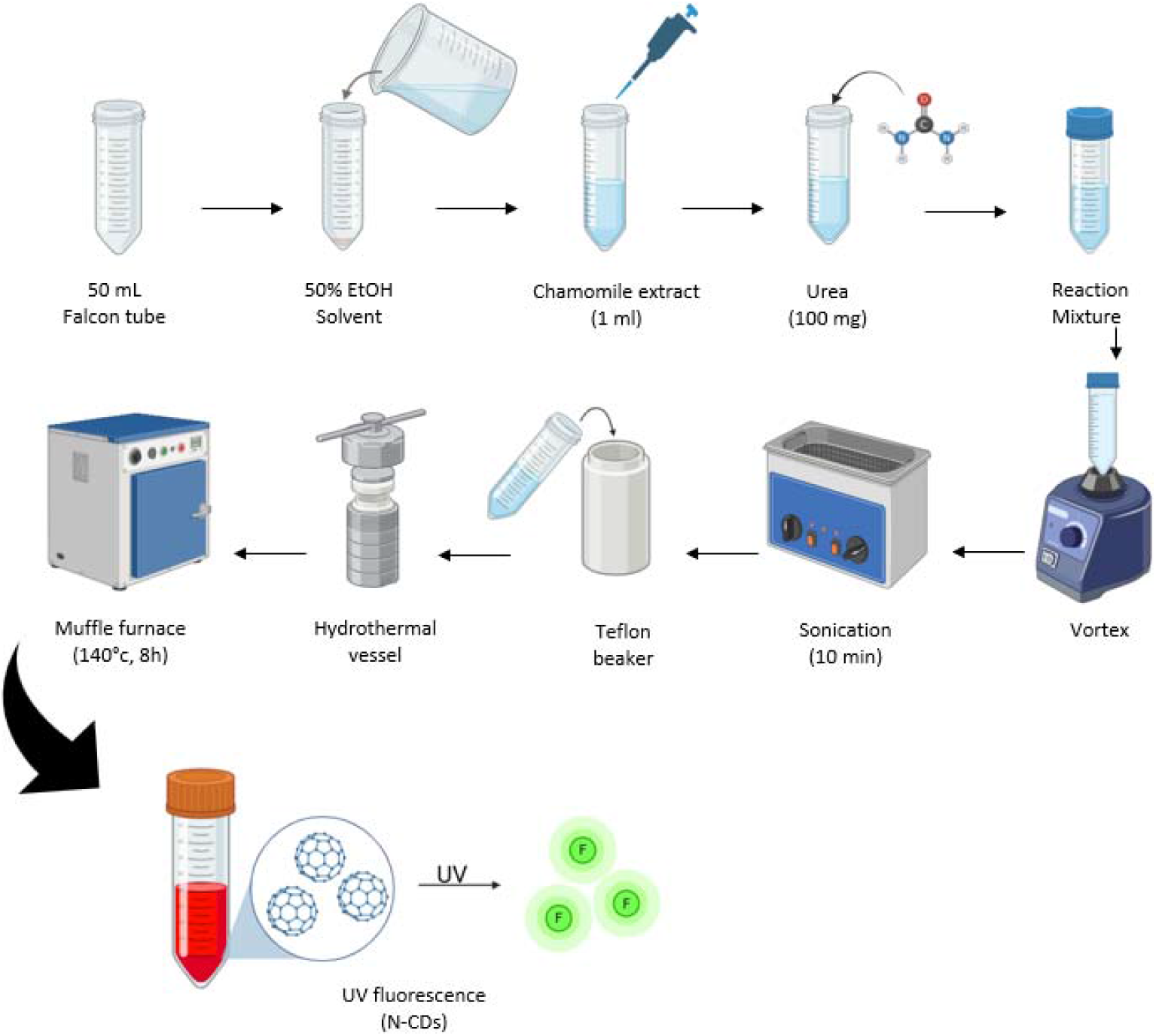
Schematic representation of the solvothermal green synthesis of nitrogen-doped chamomile-derived carbon dots CM ST-N) showing green fluorescence emission under UV illumination.

**Figure. 2.**
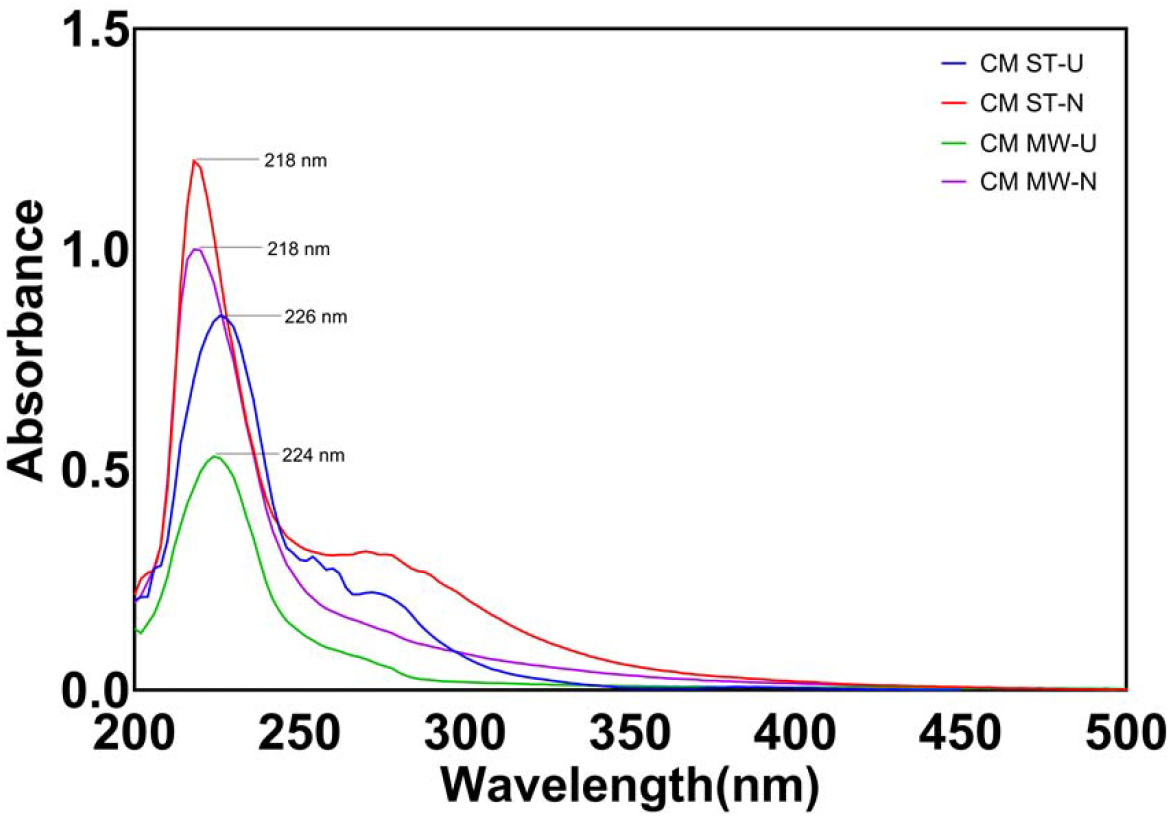
UV–visible absorption spectra of CM ST-U, CM ST-N, CM MW-U, and CM MW-N carbon dots showing characteristic π→π* and n→π* transition bands.

Comparative analysis revealed that solvothermally synthesized samples produced sharper, more intense absorption peaks relative to the microwave-assisted counterparts, suggesting a higher degree of graphitic ordering and more uniform particle formation under prolonged thermal treatment at 140°C.^33,48^ Furthermore, nitrogen-doped variants exhibited marginally elevated absorption intensities compared to their undoped equivalents, consistent with nitrogen-induced modification of the electronic structure and creation of new absorption-active surface states.^12,19^ These differences in UV–visible profiles reflect the combined influence of synthesis route and heteroatom incorporation on the optical characteristics of the resulting carbon dots, and are corroborated by subsequent photoluminescence analysis.

### 3.3 Photoluminescence Analysis

Photoluminescence emission spectra recorded across excitation wavelengths of 220–450 nm demonstrated excitation-dependent fluorescence behavior in all four variants (Fig. 3), a hallmark optical characteristic of carbon dots attributable to the heterogeneous distribution of emissive surface states and the polydisperse nature of their graphitic domains.^15,16^ Upon excitation at 400 nm, emission maxima were observed in the 500–520 nm spectral region across all samples, with peak positions remaining broadly consistent between variants, suggesting that the fundamental emissive chromophoric states are preserved irrespective of synthesis route or nitrogen doping status. However, emission intensity varied substantially between the four samples, revealing clear hierarchical differences in fluorescence efficiency.

**Figure. 3.**
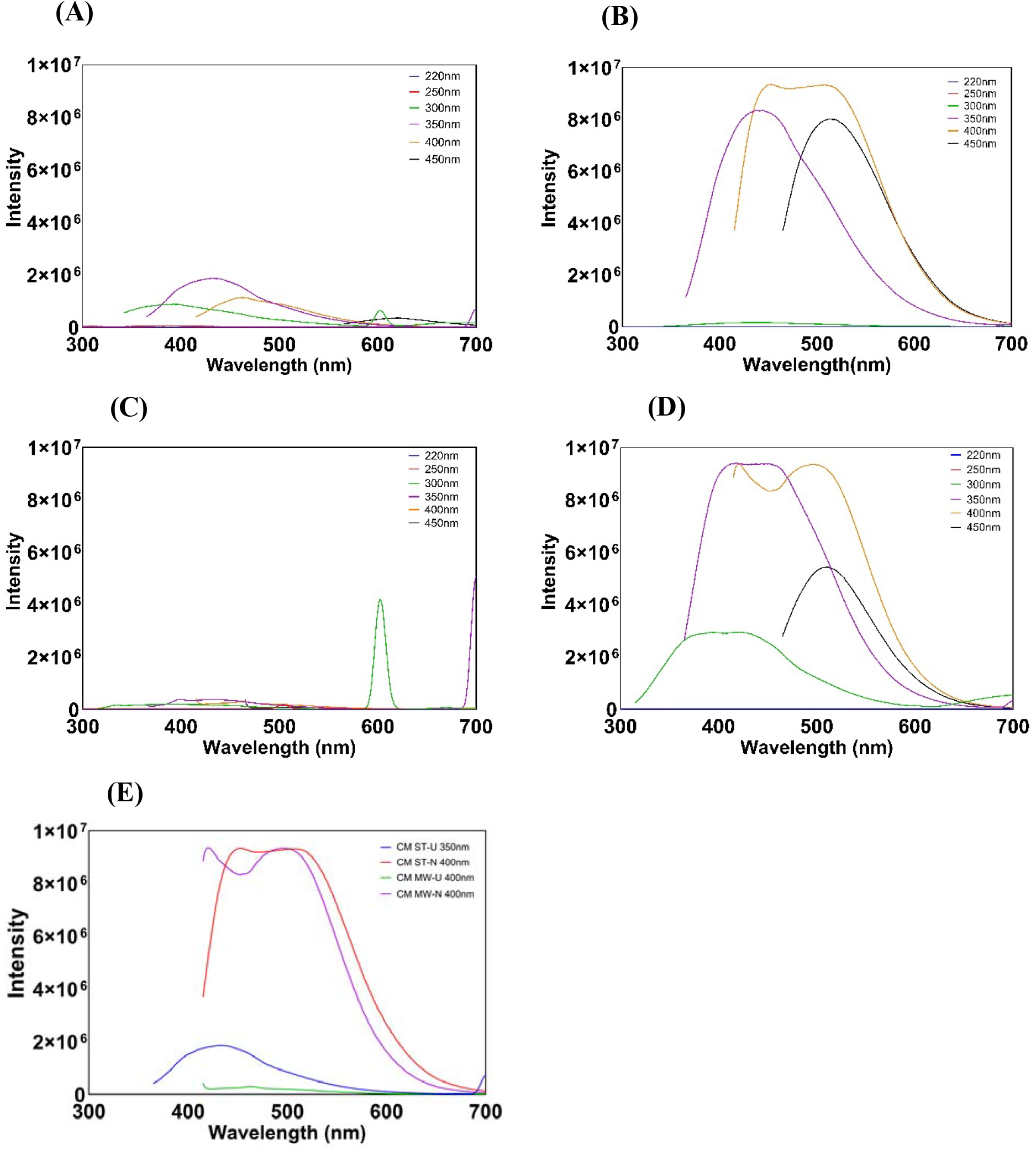
Excitation-dependent Photoluminescence Emission spectra of carbon dots. **(A)** CM ST-U, **(B)** CM ST-N, **(C)** CM MW-U, **(D)** CM MW-N, **(E)** Comparative emission intensity at optimal excitation wavelength.

CM ST-N exhibited the highest emission intensity among all variants, with a sharp, well-defined emission band and minimal spectral broadening, indicating a more uniform population of surface emissive states.^21^ CM ST-U produced comparatively weaker emission, confirming that nitrogen doping substantially enhances fluorescence efficiency in solvothermally prepared chamomile-derived CDs, consistent with previous reports attributing this enhancement to nitrogen-induced creation of additional radiative recombination pathways and improved surface passivation.^22,23^ The microwave-assisted samples—CM MW-U and CM MW-N—produced broader, less-defined emission peaks with significantly diminished intensities relative to both solvothermal variants. This reduced optical performance is ascribed to the kinetically governed, less uniform heating profile of microwave irradiation, which limits the degree of carbonization and yields particles with a more heterogeneous distribution of surface defect states, thereby reducing overall quantum efficiency.^44,49^ The superior emission characteristics of CM ST-N positioned it as the most suitable candidate for bioimaging applications and justified its selection for all subsequent detailed characterization.

### 3.4 FTIR Spectroscopic Analysis

FTIR spectroscopy was employed to identify the surface functional groups present on the synthesized carbon dots and to assess the influence of nitrogen doping on surface chemical composition (Fig. 4). All four variants displayed absorption features characteristic of functionalized carbonaceous nanomaterials, with notable differences in band intensities between nitrogen-doped and undoped samples providing direct spectroscopic evidence of heteroatom incorporation.

**Figure. 4.**
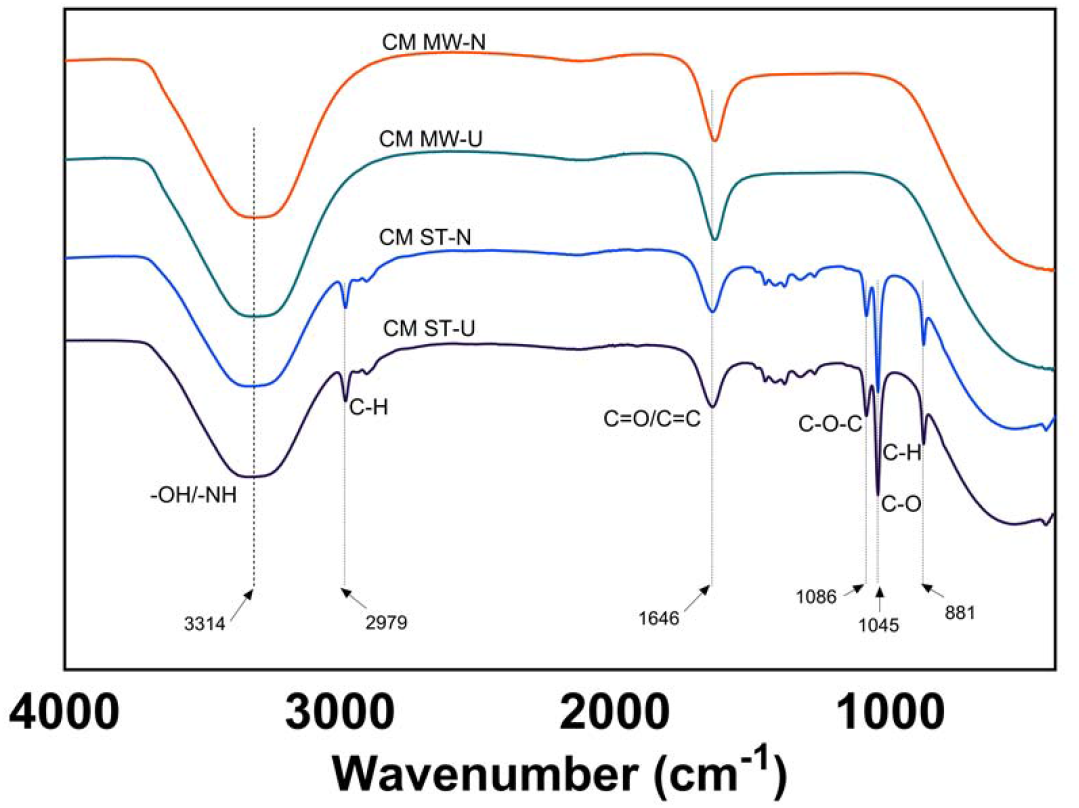
Comparative FTIR spectra of CM ST-U, CM ST-N, CM MW-U, and CM MW-N carbon dots displaying characteristic surface functional group absorption bands.

A broad, prominent absorption band centered at approximately 3314 cm^−1^ was observed across all samples, assignable to overlapping O–H and N–H stretching vibrations arising from surface hydroxyl and amine functionalities.^47,50^ This band was markedly more intense in the nitrogen-doped variants, particularly CM ST-N, providing direct spectroscopic evidence that urea-derived nitrogen-containing groups were successfully introduced onto the carbon dot surface during the solvothermal doping process. A peak near 2979 cm^−1^ present in all samples was attributed to aliphatic C–H stretching, reflecting the organic character of the chamomile precursor retained in the carbon framework. The strong absorption band at approximately 1646 cm^−1^ was assigned to C=O stretching and C=C skeletal vibrations, indicative of carbonyl functionalities and graphitic conjugated domains; the greater prominence of this band in solvothermally prepared samples is consistent with more extensive carbonization and development of conjugated π-networks under sustained heating at 140°C.^35,37^ Peaks observed at 1086 cm^−1^ and 1045 cm^−1^ were assigned to C–O–C and C–O stretching modes from ether and alcohol surface groups, respectively, while a minor absorption feature at 881 cm^−1^ was attributed to aromatic C–H bending, confirming the presence of a functionalized aromatic carbon framework.^18,47^he FTIR profile of CM ST-N—characterized by the most intense O–H/N– H, C=O, and C–O related bands among all variants—reflects the richest and most diverse surface functionalization, encompassing hydroxyl, carboxyl, carbonyl, ether, and amine moieties. This surface chemical richness is particularly advantageous for bioimaging applications: hydroxyl and carboxyl groups confer aqueous dispersibility and colloidal stability under physiological conditions, while amine functionalities provide reactive handles for potential bioconjugation and contribute directly to enhanced photoluminescence through surface state passivation.^19,20^ The spectroscopic data are entirely consistent with previous reports on nitrogen-doped carbon dots synthesized from biomass precursors via hydrothermal and solvothermal routes.^22,23^

### 3.5 Dynamic Light Scattering and Zeta Potential Analysis

DLS measurements of CM ST-N revealed a narrow, unimodal particle size distribution with a dominant peak at approximately 8.3 nm hydrodynamic diameter (Fig. 5), confirming the formation of nanoscale particles within the dimensional range canonically associated with carbon dots.^51,52^ The sharpness and unimodality of the size distribution peak, with an absence of secondary peaks at larger size ranges, indicate a highly uniform particle population with minimal inter-particle aggregation in aqueous dispersion—a characteristic of direct relevance to bioimaging, where polydisperse or aggregated probes produce heterogeneous fluorescence signals and impaired cellular uptake profiles.^7,53^

**Figure. 5.**
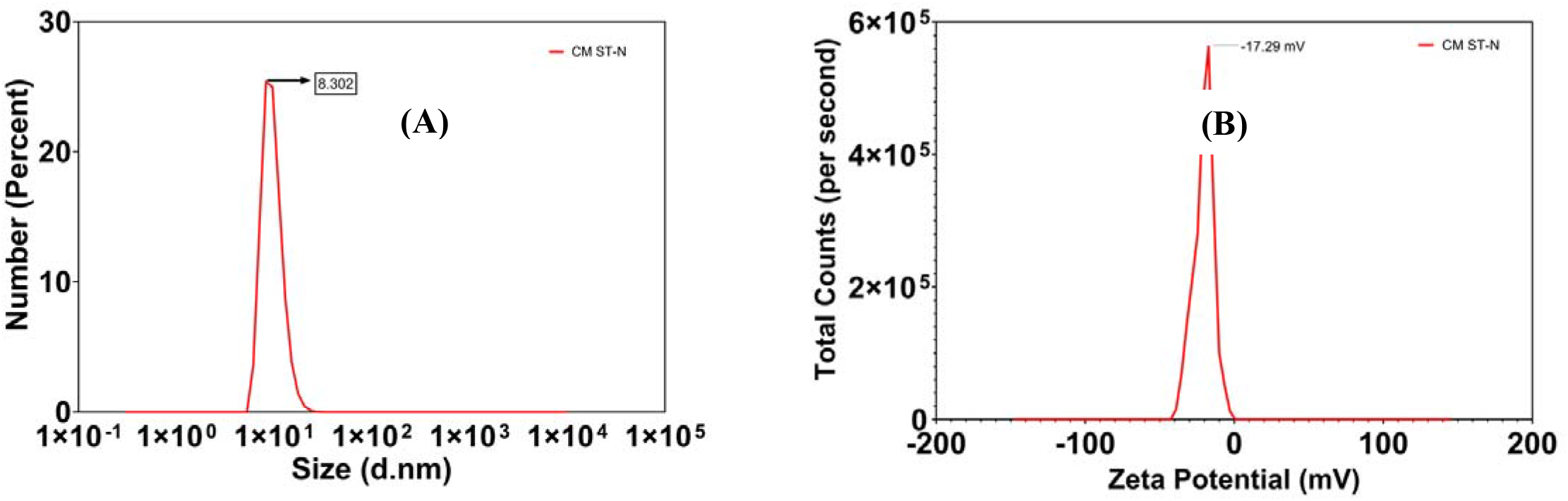
(A) DLS size distribution of CM ST-N showing a dominant peak at ~8.3 nm and (B) zeta potential analysis revealing a surface charge of −17.3 mV.

**Figure. 6.**
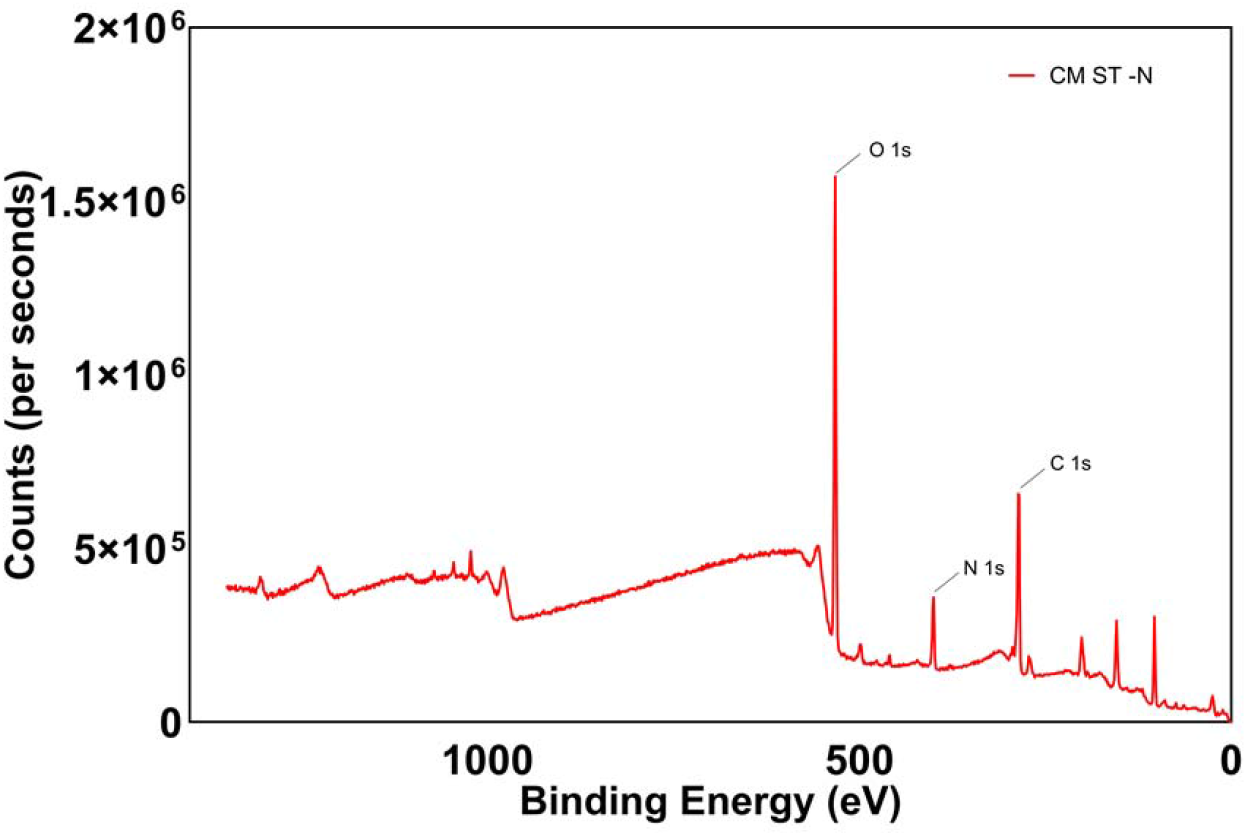
XPS survey spectrum of CM ST-N confirming elemental composition of C: 48.06%, N: 10.86%, and O: 41.08% with characteristic C 1s, N 1s, and O 1s signals.

Zeta potential analysis yielded a value of approximately −17.3 mV, reflecting the presence of negatively charged surface functional groups—primarily carboxylate and hydroxyl moieties identified by FTIR—on the carbon dot surface.^52^ While this value falls below the conventional threshold of ±30 mV typically associated with high colloidal stability, the moderate electrostatic repulsion it confers is sufficient to maintain dispersion stability in low-ionic-strength aqueous environments and is consistent with values reported for green-synthesized, surface-functionalized carbon nanodots.^22,23^ It is noteworthy that colloidal stability in biologically relevant media is governed not solely by electrostatic repulsion but also by steric stabilization arising from surface-adsorbed biomolecular species, suggesting that the functional group-rich surface of CM ST-N may provide additional steric contributions to dispersion stability in complex physiological matrices.^46^

### 3.6 X-ray Photoelectron Spectroscopy

XPS survey spectrum analysis of CM ST-N confirmed the presence of three primary elemental constituents: carbon (C 1s, ~285 eV), nitrogen (N 1s, ~399 eV), and oxygen (O 1s, ~531 eV) (Fig. 5).^50,54^ Quantitative elemental analysis yielded atomic compositions of 48.06% carbon, 10.86% nitrogen, and 41.08% oxygen. The unambiguous detection of the N 1s signal at ~399 eV provides definitive confirmation that nitrogen was successfully incorporated into the carbon dot framework during solvothermal treatment with urea, consistent with N 1s binding energies characteristic of pyrrolic N (400.0–400.5 eV) and pyridinic N (398.5–399.5 eV) configurations commonly reported for urea-doped carbon dots.^19,21^

The nitrogen content of 10.86 % achieved in CM ST-N is notably higher than values typically reported for hydrothermally synthesized nitrogen-doped CDs from biomass precursors, which generally range from 2 to 8 %.^22,23^ This elevated nitrogen incorporation likely reflects the combined contribution of urea-derived nitrogen and the endogenous nitrogen present in chamomile’s phytochemical constituents, including amino acids and nitrogen-bearing flavonoids.^35,36^ The high oxygen content (41.08 %) is consistent with the surface-oxidized nature of the carbon dots, confirming the abundance of oxygen-containing functional groups identified by FTIR, which collectively contribute to aqueous dispersibility, surface reactivity, and photoluminescence behavior.^35,47^ The XPS data thus provide quantitative validation of both the chemical composition and the successful nitrogen doping strategy, reinforcing the suitability of CM ST-N for applications requiring surface-functionalized, fluorescent carbon nanomaterials.

### 3.7 Atomic Force Microscopy

AFM topographical imaging of CM ST-N at 2 µm and 500 nm scan areas revealed the surface morphology and nanoscale spatial distribution of the synthesized particles (Fig. 7). At the larger scan scale, carbon dot features were observed as individual bright spots distributed uniformly across the mica substrate, with no evidence of microscale aggregation or cluster formation—indicative of successful dispersion and limited inter-particle interaction during sample deposition. Examination at the 500 nm scan scale resolved individual nanodot structures with height dimensions in the few-nanometer range, consistent with the hydrodynamic diameter of approximately 8.3 nm measured by DLS, noting that the AFM-derived height dimensions are expected to be smaller than the DLS hydrodynamic diameter le to the absence of the hydration shell in th d state.^45,51^

**Figure. 7.**
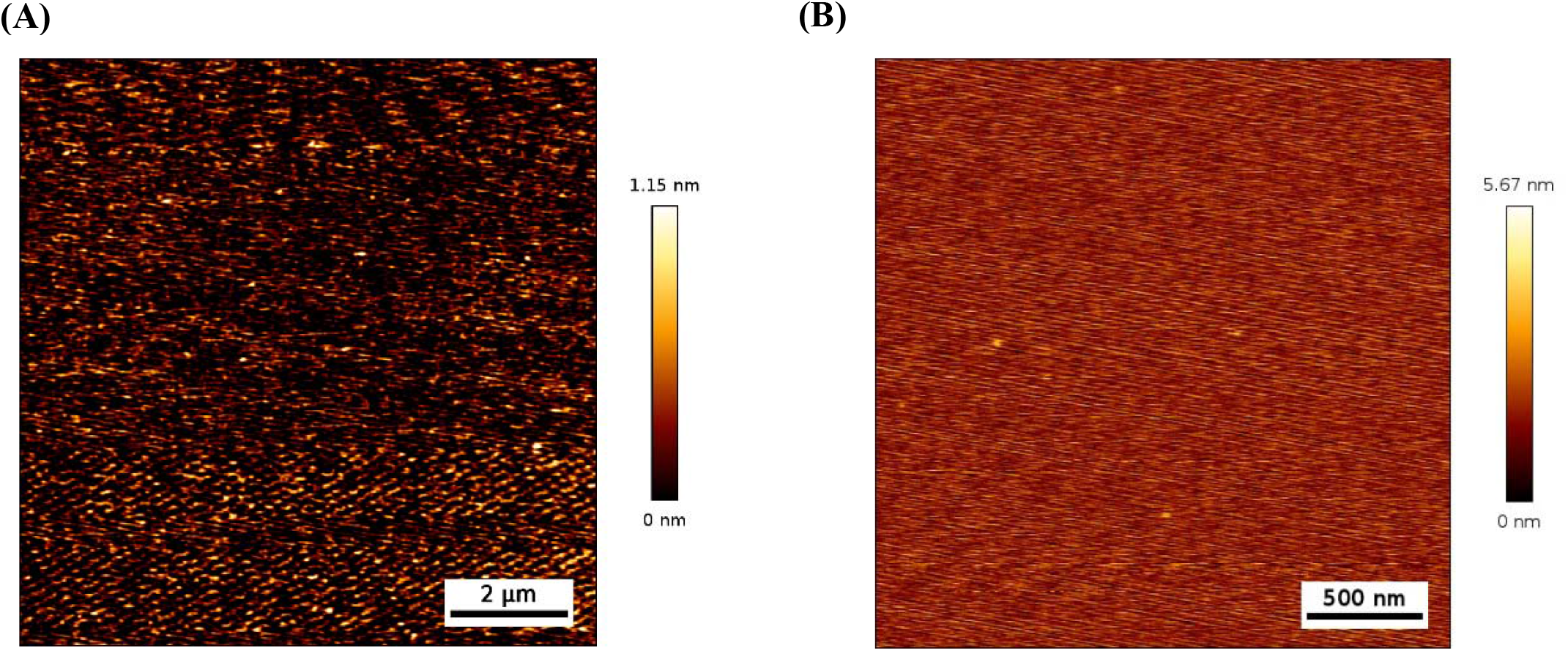
AFM topographical images of CM ST-N carbon dots at (A) 2 µm and (B) 500 nm scan areas showing uniform nanoscale distribution and absence of aggregation.

The uniform distribution and absence of aggregation across the imaged area are physically consequential for bioimaging performance, as particle aggregation is known to induce concentration-dependent fluorescence quenching, impair cellular membrane interactions, and alter endocytic uptake mechanisms relative to individually dispersed nanoprobes.^7,53^ The morphological data thus corroborate the DLS and zeta potential findings in establishing CM ST-N as a well-dispersed, nanoscale colloidal system suitable for fluorescence imaging applications. The close quantitative agreement between AFM height measurements and DLS size data further validates the structural reliability of the solvothermal synthesis approach.

### 3.8 Quantum Yield Determination

The fluorescence quantum yield of CM ST-N was determined by the comparative method using quinine sulfate (Φ = 0.54 in 0.1 M H_2_SO_4_) as the reference standard. Integrated photoluminescence intensities were plotted against absorbance values across the range 0.04– 0.07 OD for both quinine sulfate and CM ST-N, yielding linear relationships (R^2^ > 0.98) that confirm operation within the linear optical regime and the absence of significant inner filter effects. Area under the curve (AUC) analysis yielded integrated emission values of 43,214 and 43,797 for quinine sulfate and CM ST-N, respectively (Fig. 8). Application of the relative quantum yield equation yielded Φ*CM ST-N* = 0.572, corresponding to a quantum yield of 57.2%.

**Figure. 8.**
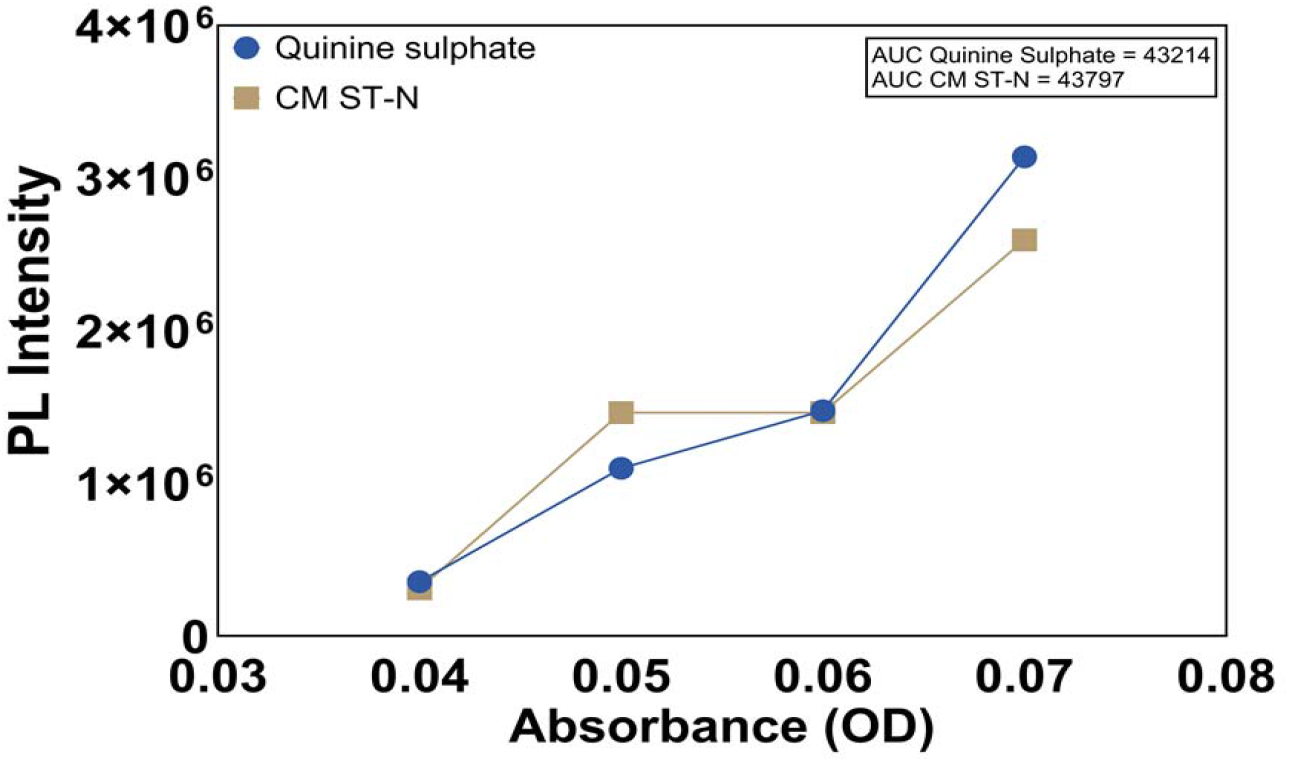
Comparative plot of integrated photoluminescence intensity versus absorbance for CM ST-N and quinine sulfate reference standard yielding a quantum yield of Φ = 57.2%.

This value represents a substantial fluorescence efficiency for a biomass-derived, nitrogen-doped carbon dot system. Comparative contextualization with the existing literature reveals that most plant-extract-derived CDs exhibit quantum yields in the range of 5–30%, with nitrogen-doped variants typically achieving 15–40%.^23,27,29^ The 57.2% quantum yield of CM ST-N therefore places it among the higher-performing biomass-derived N-CDs reported to date, attributable to the synergistic effects of effective surface passivation by chamomile phytochemical constituents, successful nitrogen incorporation creating new radiative recombination pathways, and the controlled carbonization conditions of the solvothermal route that favor the development of well-ordered emissive surface states.^22,25^ From a bioimaging perspective, high quantum yield is a primary determinant of probe sensitivity, enabling fluorescence detection at lower probe concentrations and reducing the risk of concentration-dependent cytotoxicity—a direct practical advantage of CM ST-N over lower-yield alternatives.

### 3.9 Photoluminescence Stability in Biologically Relevant Media

The photostability of a fluorescent nanoprobe in the complex chemical environment of biological media constitutes a critical prerequisite for reliable bioimaging performance.^40,43^ To assess this parameter systematically, the PL emission intensity of CM ST-N was monitored at 24, 48, and 72 hours across seven distinct solvent and media systems: Milli-Q water, methanol, absolute ethanol, 50% ethanol, PBS, DMEM, and SFM (Fig. 9).

**Figure. 9.**
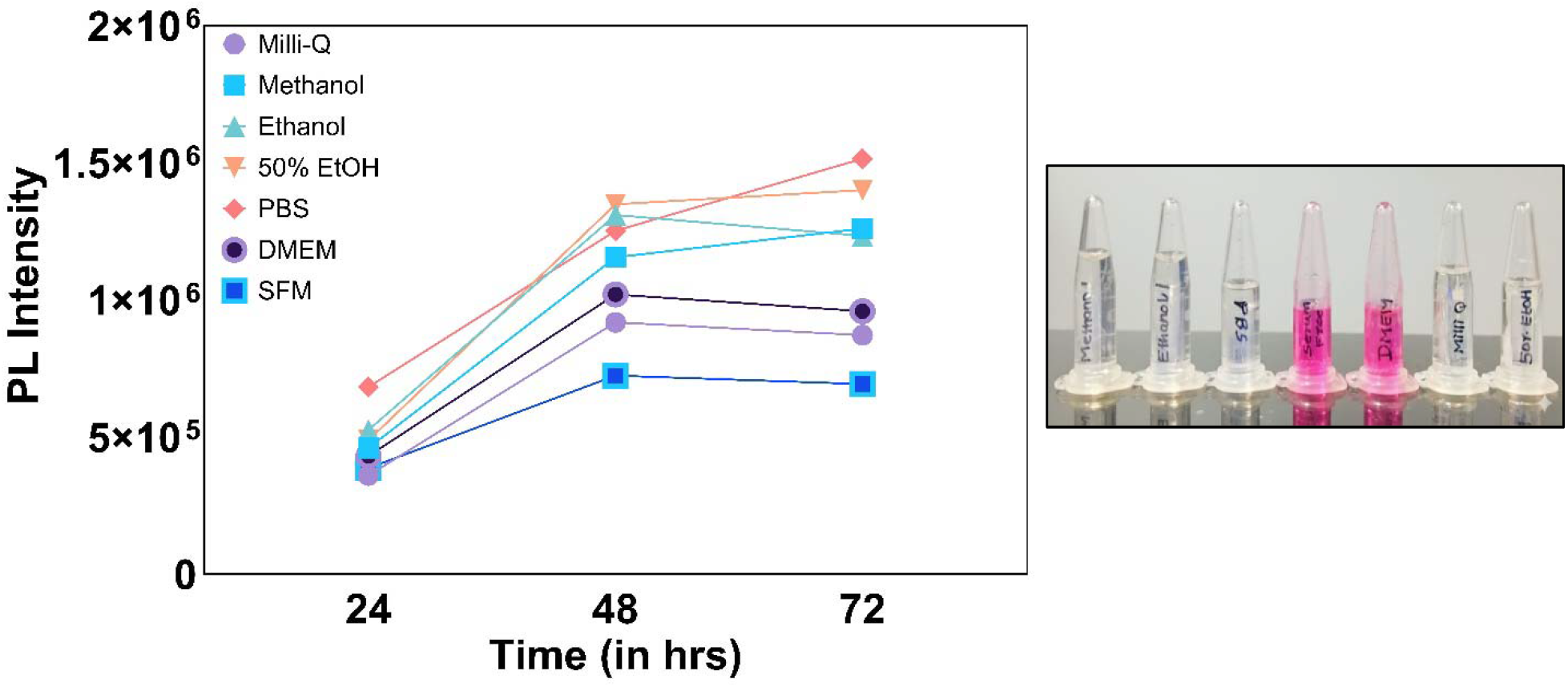
Time-dependent photoluminescence stability of CM ST-N carbon dots in Milli-Q water, methanol, ethanol, 50% ethanol, PBS, DMEM, and SFM over 72 hours.

Across all media tested, CM ST-N demonstrated sustained photoluminescence with no catastrophic loss of emission intensity over the 72-hour observation period. In PBS—the physiological ionic strength buffer most closely mimicking extracellular conditions— fluorescence intensity remained essentially stable throughout the measurement window, demonstrating that the ionic environment of a biological buffer does not adversely perturb the surface emissive states of CM ST-N. In DMEM and SFM, both of which contain complex mixtures of amino acids, vitamins, glucose, and inorganic salts relevant to cell culture conditions, initial minor fluctuations in emission intensity at the 24-hour time point were observed, followed by stabilization at consistent emission levels by 48–72 hours.^21^ This stabilization behavior is interpreted as reflecting surface equilibration of CM ST-N with media components, including potential adsorption of biomolecular species that modulate but do not ablate the surface emissive states—a phenomenon consistent with reports on green-derived CDs in complex biological matrices.^23,24^

In organic solvent media (methanol, ethanol, 50% ethanol), greater initial variability in emission intensity was observed relative to aqueous media, with intensity values converging toward stable levels by 72 hours. The absence of sustained fluorescence quenching or irreversible bleaching in any of the tested media, including conditions of high ionic strength (PBS), complex nutrient composition (DMEM, SFM), and organic solvent exposure, collectively demonstrates the robust photostability of CM ST-N across a wide range of chemical environments. This multi-media stability profile directly supports the applicability of CM ST-N as a fluorescent probe in cell culture-based bioimaging experiments, where exposure to complete growth media is unavoidable and probe stability over the duration of imaging experiments (typically 24–72 hours) is essential for reliable quantification.^2,40^

### 3.10 In Vitro Cytotoxicity Assessment

The biocompatibility of CM ST-N was rigorously evaluated using the MTT metabolic activity assay on RPE-1 retinal pigment epithelial cells across a concentration range spanning two orders of magnitude (10–500 µg mL^−1^) and three exposure durations (12, 24, and 48 hours) (Fig. 10). RPE-1 cells represent a non-cancerous, human-derived normal cell line widely employed as a standard model for cytotoxicity screening of nanomaterials intended for biomedical applications.^43,55^

**Figure. 10.**
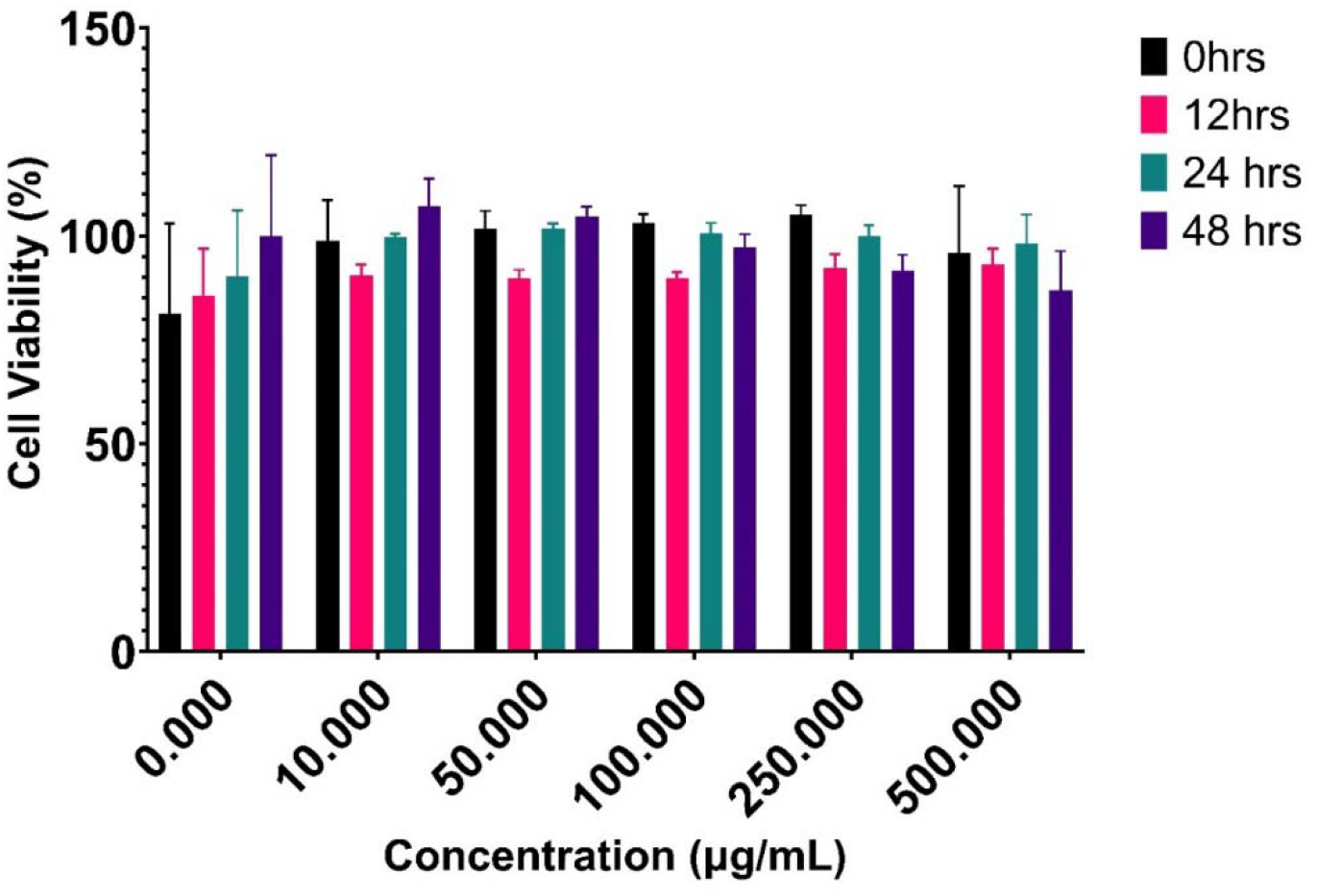
Concentration- and time-dependent cell viability of RPE-1 cells following exposure to CM ST-N carbon dots at 10–500 µg mL^−1^ for 12, 24, and 48 hours assessed by MTT assay.

At the 12-hour time point, a modest reduction in cell viability relative to untreated controls was observed across most concentrations tested. Such early-phase viability reductions are a commonly reported transient response in nanoparticle cytotoxicity studies, reflecting initial cellular adaptation to the presence of foreign particulate material rather than irreversible cytotoxic damage.^7,53^ By 24 hours, cell viability had recovered substantially across all concentrations, approaching or matching control levels at several dose points, indicating that CM ST-N does not impair mitochondrial metabolic function under sustained exposure—the mechanistic basis of the MTT assay readout. At 48 hours, a marginal reduction in viability relative to the 24-hour values was noted, consistent with the minor metabolic perturbations associated with prolonged nanomaterial exposure; critically, however, cell viability remained above 70% at all concentrations including the highest tested dose of 500 µg mL^−1^, the threshold below which a material is conventionally classified as cytotoxic according to ISO 10993-5 guidelines.

Notably, no dose-dependent reduction in cell viability was observed across the concentration range tested, indicating the absence of concentration-dependent cytotoxic mechanisms at the doses examined. This lack of dose-dependent toxicity is attributable to the physicochemical properties of CM ST-N: its small nanoscale dimensions (~8.3 nm) reduce the risk of membrane disruption, while the hydrophilic, negatively charged surface—rich in hydroxyl, carboxyl, and amine functionalities—minimizes non-specific interactions with cellular membranes and reduces cellular oxidative stress.^33,55^ The excellent biocompatibility profile of CM ST-N at concentrations up to 500 µg mL^−1^ is consistent with the broader body of literature reporting low cytotoxicity for green-synthesized, surface-functionalized carbon dots in normal human cell lines.^3,23^ The Bhatia laboratory has similarly documented the biocompatibility of carbon quantum dot systems in cellular imaging contexts, where surface chemistry and nanoscale dimensions were identified as key determinants of cytotoxic outcome.^25,34^ Taken together, the MTT data establish that CM ST-N satisfies the cytotoxic safety threshold required for deployment as a fluorescent nanoprobe in cell-based bioimaging experiments.

## 4. Conclusions

The present study reports the comparative synthesis and systematic physicochemical evaluation of four chamomile-derived carbon dot variants—CM ST-U, CM ST-N, CM MW-U, and CM MW-N—with bioimaging applicability as the primary evaluative framework. Solvothermal nitrogen doping emerged unequivocally as the superior synthetic strategy, yielding CM ST-N with a fluorescence quantum yield of 57.2%—among the highest reported for biomass-derived nitrogen-doped carbon dots—alongside well-defined nanoscale dimensions (~8.3 nm), a surface chemistry rich in hydroxyl, carboxyl, carbonyl, and amine functionalities confirmed by FTIR and XPS, moderate colloidal stability (zeta potential: −17.3 mV), and uniform, non-aggregated particle morphology established by AFM. Photoluminescence stability studies across seven solvent and biological media systems demonstrated sustained fluorescence emission over 72 hours, including in PBS, DMEM, and SFM—media directly relevant to cell culture bioimaging contexts. In vitro cytotoxicity assessment confirmed high cell viability (>70%) across the full concentration range of 10–500 µg mL^−1^ at all exposure durations tested, establishing the biocompatibility prerequisite for biological probe applications.

Collectively, CM ST-N satisfies the physicochemical requirements of a high-performance fluorescent nanoprobe for bioimaging: high quantum yield enabling sensitive detection at low probe concentrations, nanoscale dimensions favorable for cellular uptake, surface functional groups supporting aqueous dispersibility and bioconjugation potential, photostability in biologically complex media, and confirmed cytocompatibility. The green synthesis approach utilizing chamomile extract as a sustainable, phytochemically rich carbon precursor further underscores the eco-friendly credentials of this platform. These findings establish a robust physicochemical foundation for future cellular uptake studies, intracellular localization investigations, and fluorescence microscopy-based bioimaging evaluations. Further work directed toward surface functionalization with targeting ligands, exploration of two-photon excitation properties, and scale-up synthesis optimization will be necessary to fully realize the translational potential of chamomile-derived nitrogen-doped carbon dots as next-generation bioimaging nanoprobes.

